# A genome wide copper-sensitized screen identifies novel regulators of mitochondrial cytochrome c oxidase activity

**DOI:** 10.1101/2020.12.31.424969

**Authors:** Natalie M. Garza, Aaron T. Griffin, Mohammad Zulkifli, Chenxi Qiu, Craig D. Kaplan, Vishal M. Gohil

## Abstract

Copper is essential for the activity and stability of cytochrome *c* oxidase (CcO), the terminal enzyme of the mitochondrial respiratory chain. Loss-of-function mutations in genes required for copper transport to CcO result in fatal human disorders. Despite the fundamental importance of copper in mitochondrial and organismal physiology, systematic characterization of genes that regulate mitochondrial copper homeostasis is lacking. To identify genes required for mitochondrial copper homeostasis, we performed a genome-wide copper-sensitized screen using DNA barcoded yeast deletion library. Our screen recovered a number of genes known to be involved in cellular copper homeostasis while revealing genes previously not linked to mitochondrial copper biology. These newly identified genes include the subunits of the adaptor protein 3 complex (AP-3) and components of the cellular pH-sensing pathway-Rim20 and Rim21, both of which are known to affect vacuolar function. We find that AP-3 and the Rim mutants impact mitochondrial CcO function by maintaining vacuolar acidity. CcO activity of these mutants could be rescued by either restoring vacuolar pH or by supplementing growth media with additional copper. Consistent with these genetic data, pharmacological inhibition of the vacuolar proton pump leads to decreased mitochondrial copper content and a concomitant decrease in CcO abundance and activity. Taken together, our study uncovered a number of novel genetic regulators of mitochondrial copper homeostasis and provided a mechanism by which vacuolar pH impacts mitochondrial respiration through copper homeostasis.

## INTRODUCTION

Copper is an essential trace metal that serves as a cofactor for a number of enzymes in various biochemical processes, including mitochondrial bioenergetics (1). For example, copper is essential for the activity of cytochrome *c* oxidase (CcO), the evolutionarily conserved enzyme of the mitochondrial respiratory chain and the main site of cellular respiration (2). CcO metalation requires transport of copper to mitochondria followed by its insertion into Cox1 and Cox2, the two copper-containing subunits of CcO (3). Genetic defects that prevent copper delivery to CcO disrupt its assembly and activity resulting in rare but fatal infantile disorders (4, 5, 6).

Intracellular trafficking of copper poses a challenge because of the high reactivity of this transition metal. Copper in an aqueous environment of the cell can generate deleterious reactive oxygen species via Fenton chemistry (7) and can inactivate other metalloproteins by mismetallation (8). Consequently, organisms must tightly control copper import and trafficking to subcellular compartments to ensure proper cuproprotein biogenesis while preventing its toxicity. Indeed, aerobic organisms have evolved highly conserved proteins to import and distribute copper to cuproenzymes in cells (9). Extracellular copper is imported by plasma membrane copper transporters and is immediately bound to metallochaperones Atx1 and Ccs1 for its delivery to different cuproenzymes residing in the Golgi and cytosol, respectively (10).

However, copper transport to the mitochondria is not well understood. A non-proteinaceous ligand, whose molecular identity remains unknown, has been proposed to transport cytosolic copper to the mitochondria (3), where it is stored in the matrix (11). This mitochondrial matrix pool of copper is the main source of copper ions that are delivered to CcO subunits in a particularly complex process requiring multiple metallochaperones and thiol reductases (3, 12, 13). Specifically, copper from the mitochondrial matrix is exported to the intermembrane space via a yet unidentified transporter, where it is inserted into the CcO subunits by metallochaperones Cox17, Sco1, and Cox11 that operate in a bucket-brigade manner (13). The copper-transporting function of metallochaperones requires disulfide reductase activities of Sco2 and Coa6, respectively (14, 15).

In addition to the mitochondria, vacuoles in yeast and vacuole-like lysosomes in higher eukaryotes have been identified as critical storage sites and regulators of cellular copper homeostasis (16-18). Copper enters the vacuole by an unknown mechanism and is proposed to be stored as Cu(II) coordinated to polyphosphate (19). Depending on the cellular requirement, vacuolar copper is reduced to Cu(I), allowing its mobilization and export through Ctr2 (20, 21). Currently, the complete set of factors regulating the intracellular distribution of copper and its transport to the mitochondria remains unknown.

Here, we sought to identify regulators of mitochondrial copper homeostasis by exploiting the copper requirement of CcO in a genome-wide screen using a barcoded yeast deletion library. Our screen was motivated by prior observations that respiratory growth of yeast mutants such as *coa6Δ* can be rescued by copper supplementation in the media (22-24). Thus, we designed a copper-sensitized screen to identify yeast mutants whose growth can be rescued by addition of copper in the media. Our screen recovered Coa6 and other genes with known roles in copper metabolism while uncovering genes involved in vacuolar function as regulators of mitochondrial copper homeostasis. Here, we have highlighted the roles of two cellular pathways - adaptor protein 3 complex (AP-3) and the pH-sensing pathway Rim101 – that converge on vacuolar function, as important factors regulating CcO biogenesis by maintaining mitochondrial copper homeostasis.

## RESULTS

### A genome-wide copper-sensitized screen using barcoded yeast deletion mutant library

We chose the yeast, *Saccharomyces cerevisiae*, to screen for genes that impact mitochondrial copper homeostasis because it can tolerate mutations that inactivate mitochondrial respiration by surviving on glycolysis. This enables the discovery of novel regulators of mitochondrial copper metabolism whose knockout is expected to result in a defect in aerobic energy generation (25). Yeast cultured in glucose-containing media (YPD) uses glycolytic fermentation as the primary source for cellular energy, however in glycerol/ethanol-containing non-fermentable media (YPGE), yeast must utilize the mitochondrial respiratory chain and its terminal cuproenzyme, CcO, for energy production. Based on the nutrient-dependent utilization of different energy-generating pathways, we expect that deletion of genes required for respiratory growth will specifically reduce growth in non-fermentable (YPGE) medium but will not impair growth of those mutants in fermentable (YPD) medium. Moreover, if respiratory deficiency in yeast mutants is caused by defective copper delivery to mitochondria, then these mutants may be amenable to rescue via copper supplementation in YPGE respiratory growth media (Fig. 1). Therefore, to identify genes required for copper-dependent respiratory growth, we cultured the yeast deletion mutants in YPD and YPGE with or without 5 μM CuCl_2_ supplementation (Fig. 1). Our genome-wide yeast deletion mutant library was derived from the variomics library reported previously (26). It is composed of viable haploid yeast mutants, where each mutant has one nonessential gene replaced with the selection marker *KANMX4* and two unique flanking sequences (Fig. 1). These flanking sequences labeled “UP” and “DN” contains universal priming sites as well as a 20-bp barcode sequence that is specific to each deletion strain. This unique barcode sequence allows for the quantification of relative abundance of individual strain within a pool of competitively grown strains by DNA barcode sequencing (27). Here, we utilized this DNA barcode sequencing approach to quantify the relative fitness of each mutant grown in YPD and YPGE ± Cu to early stationary phase (Fig. 1).

**Figure 1.**
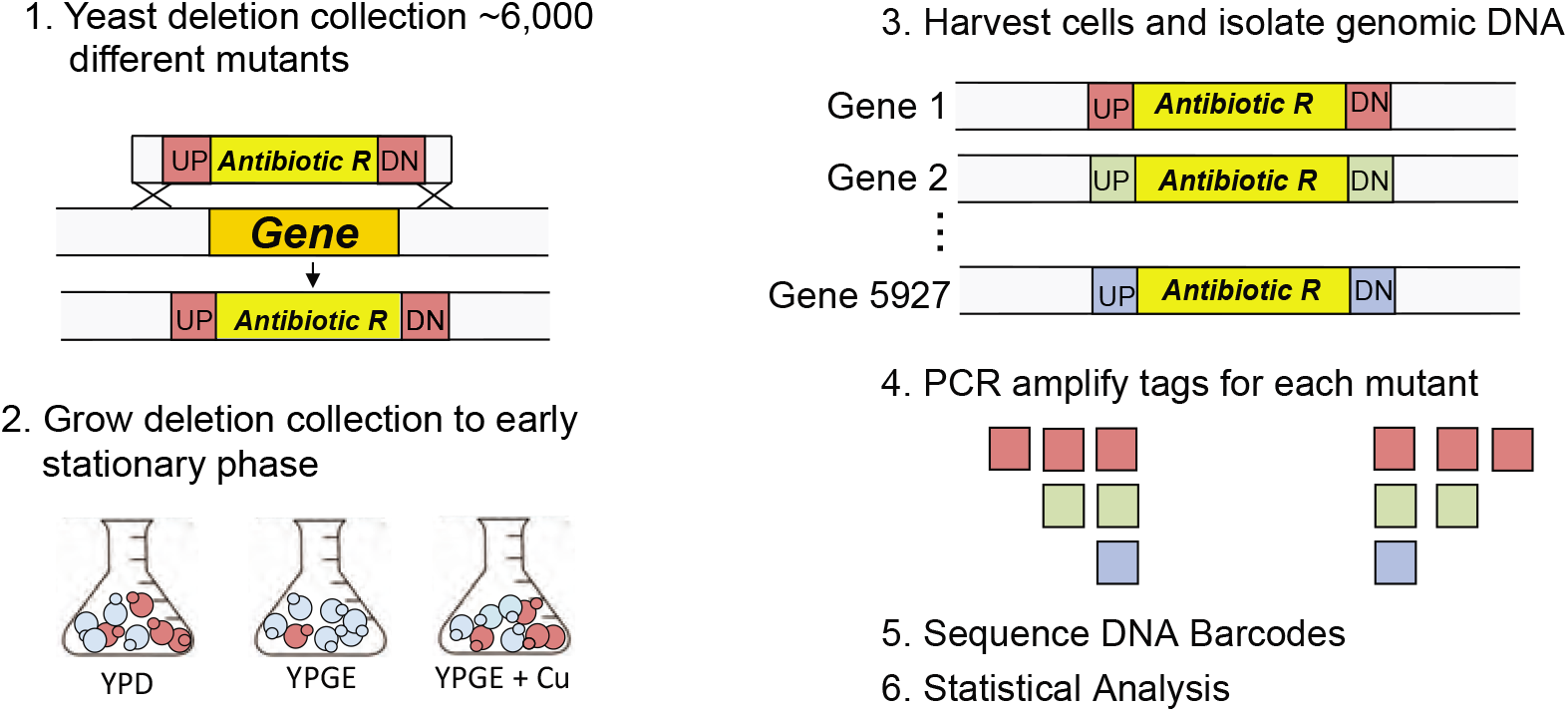
Schematic of genome-wide copper-sensitized screen. The yeast deletion library is a collection of ∼ 6000 mutants, where each mutant has a gene replaced with *kanMX4* cassette that is flanked by a unique UP tag (UP) and DOWN tag (DN) sequences. The deletion mutant pool was grown in fermentable (YPD) and non-fermentable (YPGE) medium with and without 5 μM CuCl_2_ supplementation till cells reached an optical density of 5.0. The genomic DNA was isolated from harvested cells and was used as template to amplify UP and DN tag DNA barcode sequences using universal primers. PCR products were then sequenced and the resulting data analyzed. The mutants with deletion in genes required for respiratory growth is expected to grow poorly in non-fermentable medium resulting in reduced barcode reads for that particular gene(s). However, if the same gene(s) function is supported by copper supplementation then we expect increased barcode reads for that gene(s) in copper-supplemented non-fermentable growth medium.

### Genes required for respiratory growth

We began the screen by identifying mutant strains with respiratory deficiency since perturbation of mitochondrial copper metabolism is expected to compromise aerobic energy metabolism. To identify mutants with this growth phenotype, we compared the relative abundance of each barcode in YPD to that of YPGE using T-score based on Welch’s *t*-test. T-score provides a quantitative measure of the difference in the abundance of a given mutant in two growth conditions. A negative T score identifies mutants that grow poorly in respiratory conditions; conversely, a positive T score identifies mutants with better competitive growth in respiratory conditions. We rank ordered all the mutants from negative to positive T scores and found that the lower tail of the distribution was enriched in genes with known roles in respiratory chain function as expected (Fig. 2A; Supplementary Table 1). The top “hits” representing mutants with most negative T score included *COQ3, COX5A, RCF2, COA4*, and *PET54* genes that are involved in coenzyme Q and respiratory complex IV function (Fig. 2A). To more systematically identify cellular pathways that were enriched for reduced respiratory growth, we performed gene ontology analysis using an online tool - *Gene Ontology enRichment anaLysis and visuaLizAtion* (GOrilla) (28). The gene ontology (GO) analysis identified *mitochondrial respiratory chain complex assembly* (p-value: 7.73e-23) and *cytochrome oxidase assembly* (p-value: 5.09e-22) as the top-scoring biological process categories (Fig. 2B) and *mitochondrial part* (p-value: 1.40e-25) and *mitochondrial inner membrane* (p-value: 1.48e-20) as the top-scoring molecular components category (Fig. 2C). This unbiased analysis identified the expected pathways and processes validating our screening results. We further benchmarked the performance of our screen by determining the enrichment of genes encoding for mitochondria-localized and oxidative phosphorylation (OXPHOS) proteins at three different p-value thresholds (p<0.05, p<0.025, and p<0.01) (Supplementary Fig. 1). We observed that at a p-value of <0.05, ∼25% of the genes encoded for mitochondrially localized proteins, of which ∼40% OXPHOS proteins (Supplementary Fig. 1; Supplementary Table 2). The percentage of mitochondria-localized and OXPHOS genes increased progressively as we increased the stringency of our analysis by decreasing the significance cut-off from p-value of 0.05 to 0.01 (Supplementary Fig. 1). A total of 370 genes were identified to have respiratory deficient growth at p<0.01, of which 116 are known to encode mitochondrial proteins (29), nearly half of these are OXPHOS proteins from a total of 137 known OXPHOS genes in yeast (Supplementary Fig. 1; Supplementary Table 2). Expectedly, the respiratory deficient mutants included genes required for mitochondrial NADH dehydrogenase (*NDI1*) and OXPHOS complex II, III, IV, and V as well as genes involved in cytochrome *c* and ubiquinone biogenesis, which together forms mitochondrial energy generating machinery (Fig. 2D, Supplementary Table 2). Additionally, genes encoding TCA cycle enzymes and mitochondrial DNA expression were also scored as hits (Supplementary Fig. 2). Surprisingly, a large fraction of genes required for respiratory growth encoded non-mitochondrial proteins involved in vesicle-mediated transport (Supplementary Fig. 2).

**Figure 2.**
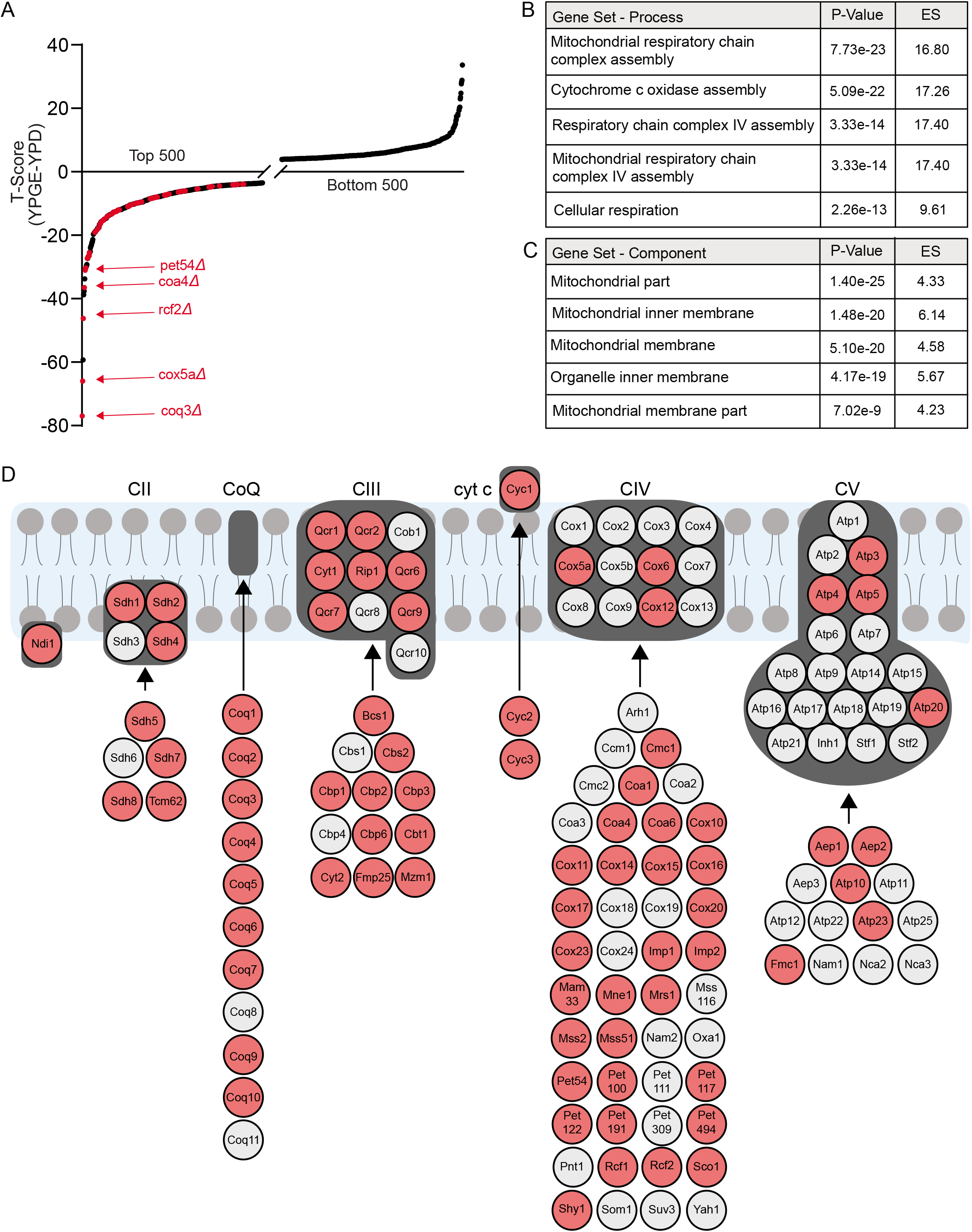
Genes required for respiratory growth. (A) Growth of each mutant in the deletion collection cultured in YPGE and YPD media was measured by BarSeq and analyzed by T-score. T(YPGE-YPD) scores are plotted for the top and bottom 500 mutants. Known mitochondrial respiratory genes are highlighted in red. (B-C) Gene ontology analysis was used to identify the top five cellular processes (B) and cellular components (C) that were significantly enriched amongst our top scoring hits from a rank ordered list, where ranking was done from the lowest to highest T-score. ES indicates enrichment score. (D) A schematic of mitochondrial OXPHOS subunits and assembly factors, where genes depicted in red were “hits” in the screen with their T-scores values below −2.35 (p-value ≤ 0.05).

### Pathway analysis for Copper-based rescue

Next, we focused on identifying mutants in which copper supplementation improved their fitness in respiratory growth conditions by comparing their abundance in YPGE + 5 μM CuCl_2_ versus YPGE growth conditions. We rank ordered the genes from positive to negative T scores. Mutants with positive T score are present in the upper tail of the distribution that displayed improved respiratory growth upon copper supplementation (Fig. 3A, Supplementary Table 3). Notably, several genes known to be involved in copper homeostasis were recovered as high scoring “hits” in our screen and were present in the expected upper tail of distribution (Fig. 3A). For example, we recovered *CTR1*, which encodes the plasma membrane copper transporter (30), *ATX1*, which encodes a metallochaperone involved in copper trafficking to the Golgi body (31), *GEF1* and *KHA1* which encodes proteins involved in copper loading into the cuproproteins in the Golgi compartment (22, 32), *GSH1* and *GSH2* which are required for biosynthesis of copper-binding molecule glutathione, and *COA6*, which encodes a mitochondrial protein that we previously discovered to have a role in copper delivery to the mitochondrial CcO (15, 23, 33) (Fig. 3A). Nevertheless, for many of our other top scoring hits, evidence supporting their role in mitochondrial copper homeostasis was either limited or lacking entirely.

**Figure 3.**
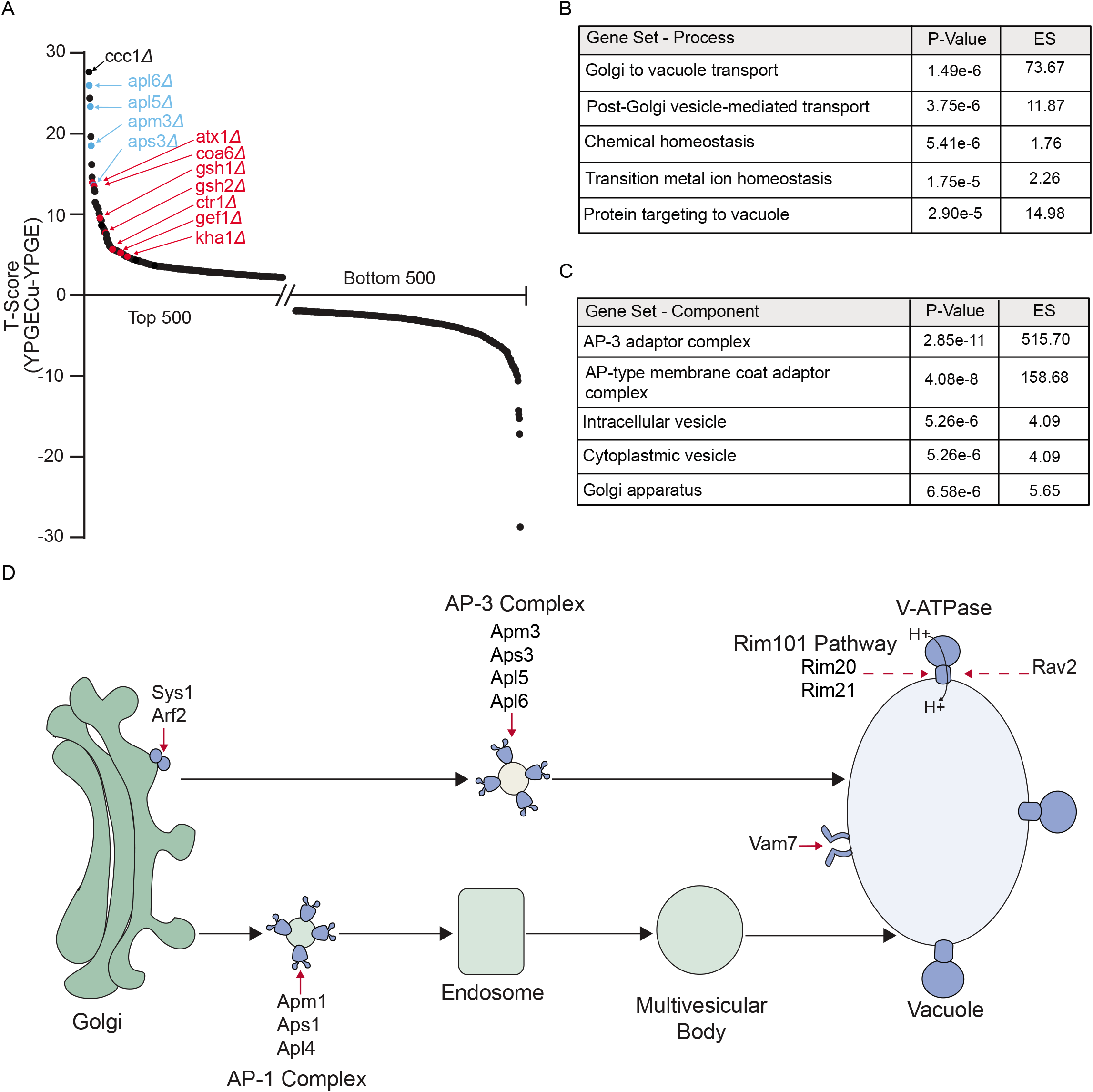
Genes required for copper homeostasis. (A) T(YPGECu-YPGE) score is plotted for top and bottom 500 mutants. Known copper homeostasis genes are highlighted in red. AP-3 subunits are highlighted in blue. (B-C) Gene ontology analysis was used to identify the top five cellular processes (B) and cellular components (C) that were significantly enriched in our top scoring hits. ES indicates enrichment score. (D) Secretory pathway mutants that displayed significantly improved growth in YPGE+Cu are displayed in blue. A dashed arrow indicates that the proteins listed are not a subunit of the complex but are involved in the maintenance of listed complex.

To determine which cellular pathways are essential for maintaining copper homeostasis, we performed gene ontology analysis using GOrilla. GO analysis identified biological processes - *Golgi to vacuole transport* (p-value: 1.49e-6), and *post-Golgi vesicle-mediated transport*, (p-value: 3.75e-6) as the most significantly enriched pathways (Fig. 3B). Additionally, GO category *transition metal ion homeostasis* - was also in the top five significantly enriched pathways, (p-value: 1.75e-5) (Fig. 3B). GO analysis for cellular component categories identified adaptor protein 3 complex (AP-3), which is known to transport vesicles from the Golgi body to vacuole, as the top scoring cellular component (p-value: 2.85e-11) (Fig. 3C). All four subunits of AP-3 complex (*APL6, APM3, APL5, APS3)* complex were in the top 10 of our rank list (Fig. 3A, Supplementary Table 3) (34, 35). Additionally, two subunits of the Rim101 pathway (*RIM20* and *RIM21)*, both of which are linked to vacuolar function (36), were also in our list of top-scoring genes (Supplementary Table 3). Of note, the seven major components of the Rim101 pathway were identified as top-scoring hits for respiratory deficient growth (Supplementary Fig. 2). Placing the hits from our screen on cellular pathways revealed a number of “hits” that were either involved in Golgi bud formation (Sys1, Arf2), vesicle coating (AP-3 and AP-1 complex subunits), tethering and fusion of Golgi vesicle cargo to the vacuole (Vam7), and vacuolar ATPase expression and assembly (Rim20, Rim21, Rav2) (Fig. 3D). We reasoned that these biological processes and cellular components were likely high scoring due to the role of the vacuole as a major storage site of intracellular metals (16). We decided to focus on AP-3 and Rim mutants, as these cellular components were not previously linked to mitochondrial respiration or mitochondrial copper homeostasis.

### AP-3 mutants exhibit reduced abundance of CcO and V-ATPase subunits

To validate our screening results and to determine the specificity of the copper-based rescue of AP-3 mutants, we compared the respiratory growth of AP-3 deletion strains, *aps3Δ, apl5Δ*, and *apl6Δ* on YPD and YPGE media with or without Cu, Mg, and Zn supplementation. Each of the AP-3 mutants exhibited reduced respiratory growth in YPGE media at 37°C, which was fully restored by copper, but not by magnesium or zinc (Fig. 4A), indicating that the primary defect in these cells is dysregulated copper homeostasis. Here we used 37°C for growth measurement to fully uncover growth defect on solid media. The *coa6Δ* mutant was used as a positive control because we have previously shown that respiratory growth deficiency of *coa6Δ* can be rescued by Cu supplementation (23). Since recent work has identified the role of the yeast vacuole in mitochondrial iron homeostasis (37, 38) we asked if iron supplementation could also rescue the respiratory growth of AP-3 mutants. Unlike copper, which rescued respiratory growth of AP-3 mutants at 5 μM concentration, low concentrations of iron (≤ 20 μM) did not rescue respiratory growth; but we did find that high iron supplementation (100 μM) improved their respiratory growth (Supplementary Fig. 3). To uncover the biochemical basis of reduced respiratory growth, we focused on Cox2, a copper-containing subunit of CcO, whose stability is dependent on copper availability and whose levels serve as a reliable proxy for mitochondrial copper content. The steady state levels of Cox2 were modestly but consistently reduced in all four AP-3 mutants tested (Fig. 4B).

**Figure 4.**
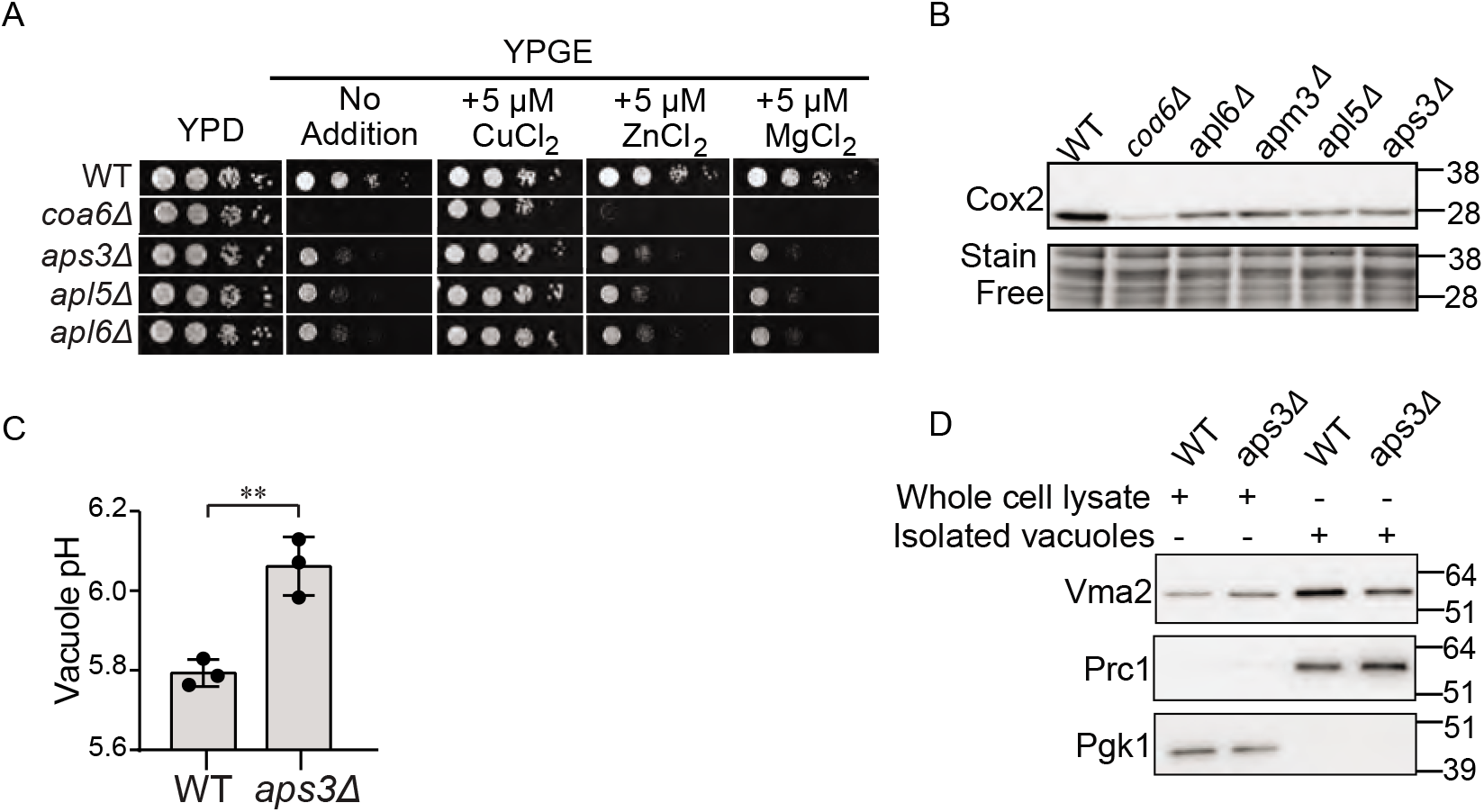
Loss of AP-3 results in reduced vacuolar and mitochondrial function. (A) Serial dilutions of WT and the indicated mutants were seeded onto YPD and YPGE plates with and without 5 μM CuCl_2_, MgCl_2_ and ZnCl_2_ and grown at 37°C for two (YPD) or four days (YPGE). *coa6Δ* cells, which have been previously shown to be rescued by CuCl_2_, were used as a control. (B) Whole cell protein lysate was analyzed by SDSPAGE/western blot using a Cox2 specific antibody to detect CcO abundance. Stain free imaging served as a loading control. *coa6Δ* cell lysate was used as control for decreased Cox2 levels. (C) Vacuolar pH of WT and *aps3Δ* cells was measured by using BCECF-AM dye. (D) Whole cell lysate and isolated vacuole fractions were analyzed by SDSPAGE/western blot. Vma2 was used to determine V-ATPase abundance. Prc1 and Pgk1 served as loading controls for vacuole and whole cell protein lysate, respectively. Data are expressed as mean ± SD; **p < 0.01, (n = 3). Each data point represents a biological replicate.

AP-3 complex function has not been directly linked to mitochondria but is linked to the trafficking of proteins from the Golgi body to the vacuole. Therefore, the decreased abundance of Cox2 in AP-3 mutants could be due to an indirect effect involving the vesicular trafficking role of the AP-3 complex. A previous study has shown that the AP-3 complex interacts with a subunit of the V-ATPase in human cells (34). As perturbation in V-ATPase function had been linked to defective respiratory growth (37-41), we wondered if AP-3 impacts mitochondrial function via trafficking V-ATPase subunit(s) to the vacuole. To test this idea, we first measured vacuolar acidification and found that the AP-3 mutant, *aps3Δ*, exhibited significantly increased vacuolar pH (Fig. 4C). We hypothesized that the elevated vacuolar pH of *aps3Δ* cells could be due to a perturbation in the trafficking of V-ATPase subunit(s). To test this possibility, we measured the levels of V-ATPase subunit Vma2, in wild type (WT) and *aps3Δ* cells, by western blotting and found that Vma2 levels were indeed reduced in the isolated vacuolar fractions of *aps3Δ* cells but were unaffected in the whole cells (Fig. 4D). The decreased abundance of Vma2 in vacuoles of yeast AP-3 mutant explains decreased vacuolar acidification because Vma2 is an essential subunit of V-ATPase. Taken together, these results suggest that the AP-3 complex is required for maintaining vacuolar acidification, which in turn could impact mitochondrial copper homeostasis.

### Genetic defects in Rim101 pathway perturbs mitochondrial copper homeostasis

Next, we focused on two other hits from the screen, Rim20 and Rim21, which are the members of the Rim101 pathway that has been previously linked to the V-ATPase expression (42-45). The loss of Rim101 results in the decreased expression of V-ATPase subunits (43, 44). Consistently, we found elevated vacuolar pH in *rim20Δ* cells (Fig. 5A). We then compared the respiratory growth of *rim20Δ* and *rim21Δ* on YPD and YPGE media with or without Cu, Mg, or Zn supplementation. Consistent with our screening results, these mutants exhibited reduced respiratory growth that was fully restored by copper but not magnesium or zinc (Fig. 5B). To directly test the roles of these genes in cellular copper homeostasis, we measured the whole-cell copper levels of *rim20Δ* by inductively coupled plasma mass spectrometry (ICP-MS). The intracellular copper levels under basal or copper-supplemented conditions in *rim20Δ* cells were comparable to the WT cells, suggesting that the copper import or sensing machinery is not defective in this mutant (Fig. 5C). In contrast to the total cellular copper levels, *rim20Δ* did exhibit significantly reduced mitochondrial copper levels, which were restored by copper supplementation (Fig. 5D).

**Figure 5.**
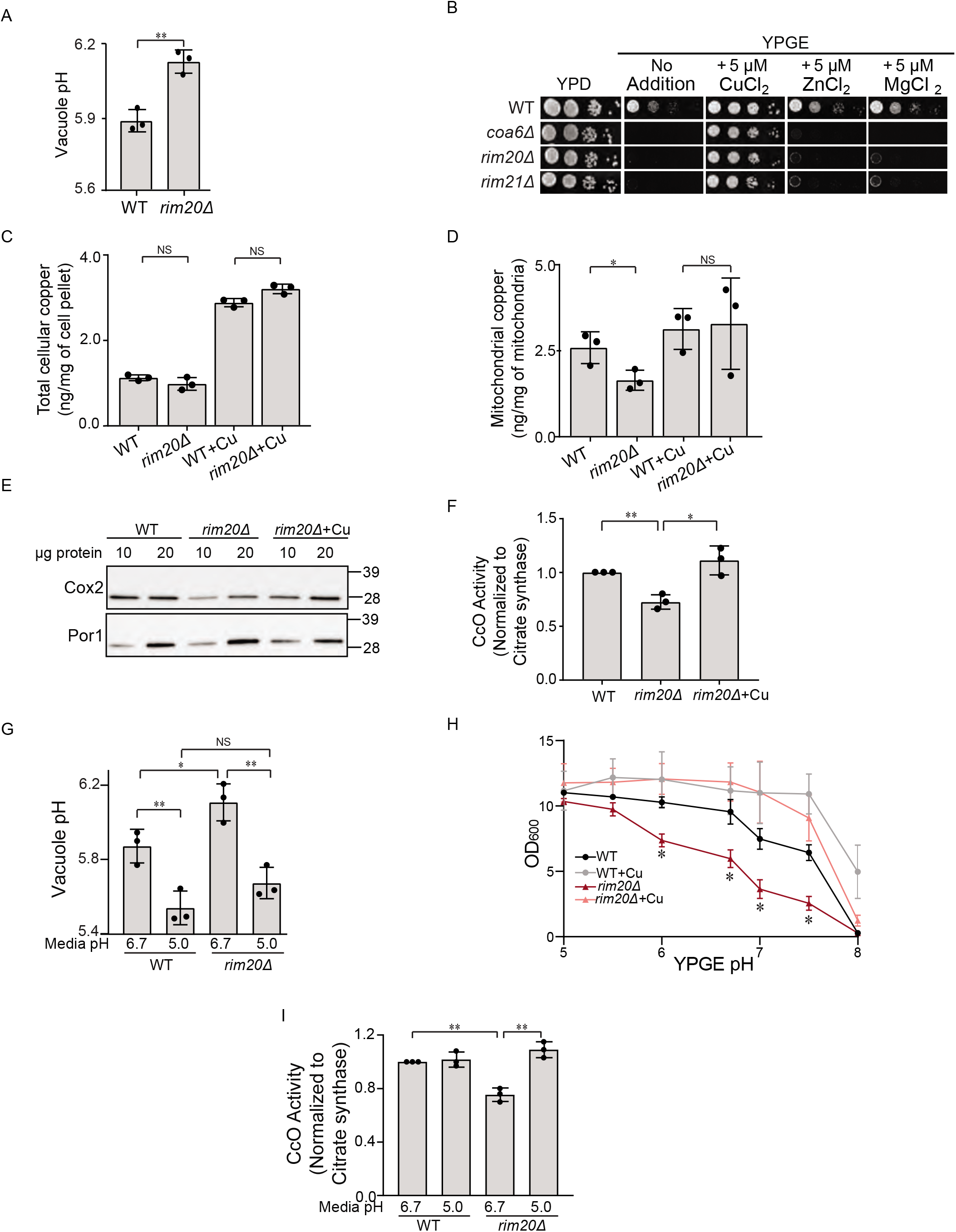
Normalization of vacuolar pH in *rim20Δ* cells restores mitochondrial copper homeostasis. (A) Vacuolar pH of WT and *rim20Δ* cells was measured by BCECF-AM dye. (B) Serial dilutions of WT and the indicated mutants were seeded onto YPD and YPGE plates with and without 5 μM CuCl_2_, MgCl_2_, or ZnCl_2_ and grown at 37°C for two (YPD) or four days (YPGE). (C) Cellular and (D) mitochondrial copper levels were measured by ICP-MS. (E) Mitochondrial proteins were analyzed by SDS-PAGE/western blot. Cox2 served as a marker for CcO levels, and Por1 served as a loading control. (F) CcO activity was measured spectrophotometrically and normalized to the citrate synthase activity. (G) Vacuolar pH of WT and *rim20Δ* cultured in standard (pH 6.7) or acidified (pH 5.0) YPGE medium was measured by BCECF-AM dye. (H) The optical density of WT and *rim20Δ* cultures after 42 hours growth in YPGE medium at the indicated pH values with or without 5 μM CuCl_2_. (I) CcO activity of WT and *rim20Δ* cultured in standard or acidified YPGE was normalized to citrate synthase activity. Data are expressed as mean ± SD; NS = not significant, *p<0.05, **p < 0.01, (n = 3). Each data point represents a biological replicate.

The decrease in mitochondrial copper levels is expected to perturb the biogenesis of CcO in *rim20Δ* cells. Therefore, we measured the abundance and activity of this complex by western blot analysis and enzymatic assay, respectively. Consistent with the decrease in mitochondrial copper levels, *rim20Δ* cells exhibited a reduction in the abundance of Cox2 along with a decrease in CcO activity, both of which were rescued by copper supplementation (Fig. 5E and F). To further dissect the compartment-specific effect by which Rim20 impacts cellular copper homeostasis, we measured the abundance and activity of Sod1, a mainly cytosolic cuproenzyme. We found that unlike CcO, Sod1 abundance and activity remain unchanged in *rim20Δ* cells (Supplementary Fig. 4).

To determine if the decrease in CcO activity in the absence of Rim20 was due to its role in maintaining vacuolar pH, we manipulated vacuolar pH by changing the pH of the growth media. Previously, it has been shown that vacuolar pH is influenced by the pH of the growth media through endocytosis (46, 47). Indeed, acidifying growth media to pH 5.0 from the basal pH of 6.7 normalized vacuolar pH of *rim20Δ* to the WT levels and both strains exhibited lower vacuolar pH when grown in acidified media (Fig. 5G). Under these conditions of reduced vacuolar pH, the respiratory growth of *rim20Δ* was restored to WT levels (Fig. 5H). Notably, alkaline media also reduced the respiratory growth of WT cells, though the extent of growth reduction was lower than *rim20Δ*, which is likely because of a fully functional V-ATPase in WT cells (Fig. 5H). The restoration of respiratory growth by copper supplementation was independent of growth media pH (Fig. 5H). To uncover the biochemical basis of the restoration of respiratory growth of *rim20Δ* by acidified media, we measured CcO enzymatic activity in WT and *rim20Δ* cells grown in either basal or acidified growth medium (pH 6.7 and 5.0), respectively. Consistent with the respiratory growth rescue, the CcO activity was also restored in cells grown at an ambient pH of 5.0 (Fig. 5I). Taken together, these findings causally links vacuolar pH to CcO activity via mitochondrial copper homeostasis.

### Pharmacological inhibition of the V-ATPase results in decreased mitochondrial copper

To directly assess the role of vacuolar pH in maintaining mitochondrial copper homeostasis, we utilized Concanamycin A (ConcA), a small molecule inhibitor of V-ATPase. Treating WT cells with increasing concentrations of ConcA led to progressively increased vacuolar pH (Fig. 6A). Notably, the increase in vacuolar pH with pharmacological inhibition of V-ATPase by ConcA was much more pronounced (Fig. 6A) than via genetic perturbation in *aps3Δ* or *rim20Δ* cells (Figs. 4C and 5A). Correspondingly, we observed a pronounced decrease in CcO abundance and activity in ConcA treated cells (Fig. 6B, C). This decrease in abundance of CcO is likely due to a reduction in mitochondrial copper levels (Fig. 6D). This data establishes the role of the vacuole in regulating mitochondrial copper homeostasis and CcO function.

**Figure 6.**
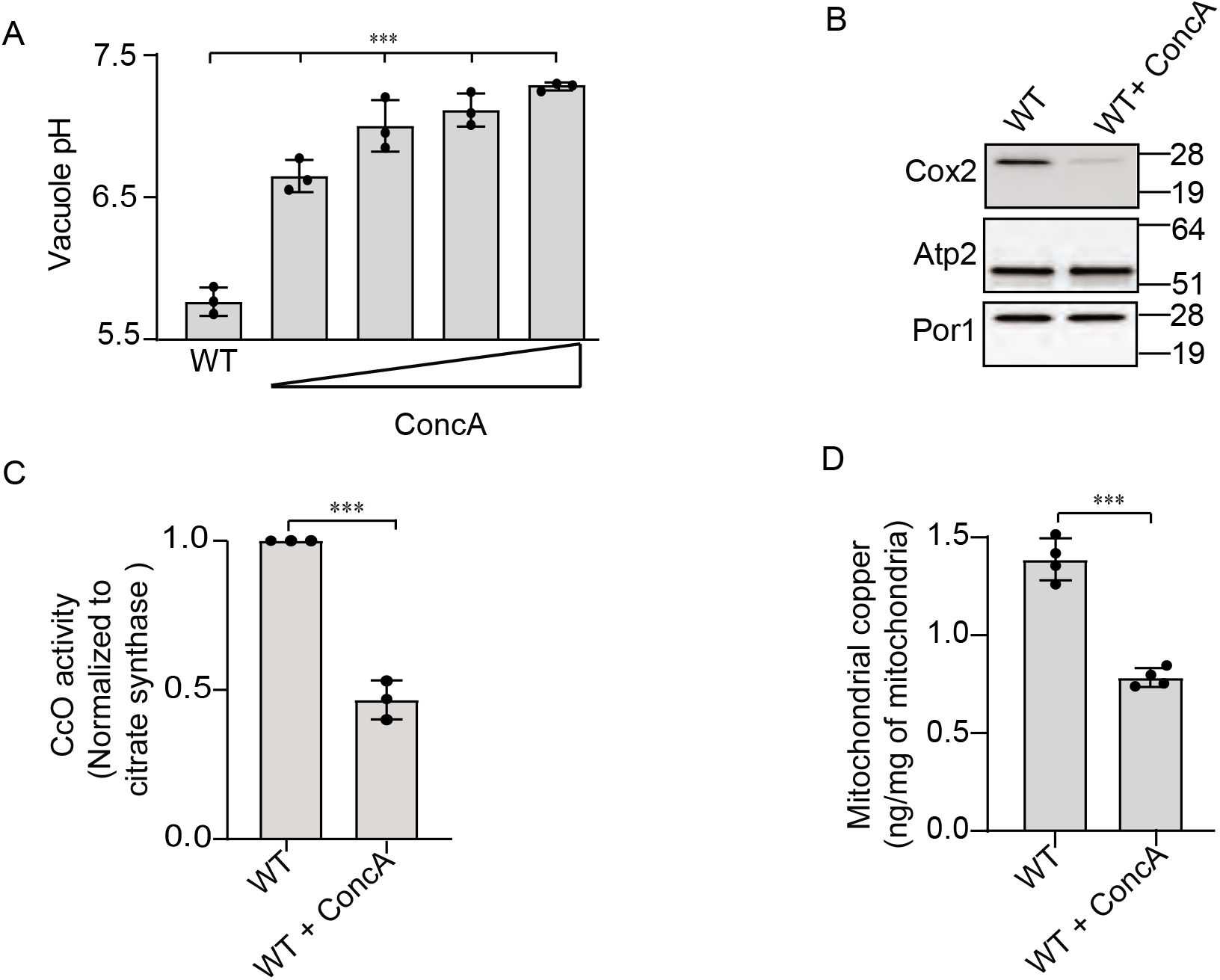
Pharmacological inhibition of V-ATPase decreases mitochondrial copper content. (A) Vacuolar pH of WT cells grown in the presence of either DMSO or 125, 250, 500, 1000 nM ConcA. (B) Mitochondrial proteins in WT cells treated with DMSO or 500 nM ConcA were analyzed by SDSPAGE/western blot. Cox2 served as a marker for CcO abundance, Atp2 and Por1 were used as loading controls. (C) CcO activity in WT cells treated with DMSO or 500 nM ConcA is shown after normalization with citrate synthase activity. (D) Mitochondria copper levels in WT cells treated with DMSO or 500 nM ConcA were determined by ICP-MS. Data are expressed as mean ± SD; ***p < 0.001, (n = 3 or 4). Each data point represents a biological replicate.

## DISCUSSION

Mitochondria are the major intracellular copper storage sites that harbor important cuproenzymes like CcO. When faced with copper deficiency, cells prioritize mitochondrial copper homeostasis suggesting its critical requirement for this organelle (48). However, the complete set of factors required for mitochondrial copper homeostasis has not been identified. Here, we report a number of novel genetic regulators of mitochondrial copper homeostasis that link mitochondrial bioenergetic function with vacuolar pH. Specifically, we show that when vacuolar pH is perturbed by genetic, environmental, or pharmacological factors, then copper availability to the mitochondria is limited, which in turn reduces CcO function and impairs aerobic growth and mitochondrial respiration.

It has been known for a long time that V-ATPase mutants have severely reduced respiratory growth (39, 40) and more recent high-throughput studies have corroborated these observations (49-51). However, the molecular mechanisms underlying this observation have remained obscure. Recent studies have shown that a decrease in vacuolar acidity (i.e. increased vacuolar pH) perturbs cellular and mitochondrial iron homeostasis, which impairs mitochondrial respiration, as iron is also required for electron transport through the mitochondrial respiratory chain due to its role in iron-sulfur cluster biogenesis and heme biosynthesis (37, 38, 52, 53). In an elegant series of experiments, Hughes et al, showed that when V-ATPase activity is compromised, there is an elevation in cytosolic amino acids because vacuoles with defective pH are unable to import and store amino acids. The resulting elevation in cytosolic amino acids, particularly cysteine, are toxic to the cells by disrupting cellular iron homeostasis and iron-dependent mitochondrial respiration (38). Although this exciting study took us a step closer to our understanding of V-ATPase-dependent mitochondrial function, the mechanism by which elevated cysteine perturbs iron homeostasis is still unclear. Since cysteine can strongly bind cuprous ions (54, 55) its sequestration in cytosol by cysteine would decrease its availability to Fet3, a multi-copper oxidase (56) required for the uptake of extracellular iron, which in turn would aggravate iron deficiency. Thus, a defect in cellular copper homeostasis could cause a secondary defect in iron homeostasis. Consistent with this idea, we observed a rescue of AP-3 mutants’ respiratory growth with high iron supplementation (Supplementary Fig. 3). Interestingly, AP-3 has also been previously linked to vacuolar cysteine homeostasis (57).

Our results showing diminished CcO activity and/or Cox2 levels in AP-3, Rim20, and ConcA-treated cells (Figs. 4B, 5E and F, 6B and C) connects vacuolar pH to mitochondrial copper biology. However, a modest decrease in CcO activity may not be sufficient to reduce respiratory growth. Therefore, it is very likely that the decreased respiratory growth we have observed is a result of a defect not only in copper but also in iron homeostasis. Consistent with this idea, previous high throughput studies reported sensitivity of AP-3 and Rim101 pathway mutants in conditions of iron deficiency and overload (58, 59). Moreover, Rim20 and Rim101 mutants have been shown to display sensitivity to copper starvation in *Cryptococcus neoformans*, an opportunistic fungal pathogen (60) and partial knockdown of Ap3s1, a subunit of AP-3 complex in zebrafish, sensitized developing melanocytes to hypopigmentation in low-copper environmental conditions (61). Thus, the Rim pathway and the AP-3 pathway is linked to copper homeostasis in multiple organisms. Our discovery of AP-3 pathway mutants and other mutants involved in the Golgi-to-vacuole transport (Fig. 3) is also consistent with a previous genome-wide study, which identified the involvement of these genes in Cu-dependent growth of yeast *Saccharomyces cerevisiae* (50), however, the biochemical mechanism(s) underlying the functional connection between the vacuole and mitochondrial CcO has not been previously elucidated. Thus, the results from our study are not only consistent with previous studies but also provide a biochemical mechanism elucidating how disruption in vacuolar pH perturbs mitochondrial respiratory function via copper-dependence of CcO. Interestingly, in both the genetic and pharmacological models of reduced V-ATPase function, mitochondrial copper levels were reduced (Fig. 5D and Fig. 6D) but were not absent, suggesting that the vacuole may only partially contribute to mitochondrial Cu homeostasis. Supporting this hypothesis, rescue of respiratory growth by copper supplementation was successful irrespective of vacuolar pH (Fig. 5 H).

The results of this study could also provide insights into mechanisms underlying the pathogenesis of human diseases associated with aberrant copper metabolism and/or decreased V-ATPase function including Alzheimer’s disease, amyotrophic lateral sclerosis (ALS), and Parkinson’s disease (62-68). Although multiple factors are known to contribute to the pathogenesis of these diseases, our study suggests disrupted mitochondrial copper homeostasis may also be an important contributing factor. In contrast to these multi-factorial diseases, pathogenic mutations in AP-3 subunits are known to cause Hermansky-Pudlak syndrome (HPS), a rare autosomal disorder, which is often associated with high morbidity (69-71). Just as in yeast, AP-3 in humans is required for the transport of vesicles to the lysosome, which is evolutionarily and functionally related to the yeast vacuole. Our study linking AP-3 to mitochondrial function suggests that decreased mitochondrial function could contribute to HPS pathology. More generally, decreased activity of V-ATPase has been linked to age-related decrease in lysosomal function (34, 72, 73) and impaired acidification of yeast vacuole has been shown to cause accelerated aging (41). Therefore, in addition to uncovering the fundamental aspects of cell biology of metal transport and distribution, our study suggests a possible role of mitochondrial dysfunction in multiple human disorders.

## METHODS

### Yeast strains and growth conditions

Individual yeast *Saccharomyces cerevisiae* mutants used in this study were obtained from Open Biosystems or were constructed by one-step gene disruption using a hygromycin cassette (74). All strains used in this study are listed in Table 1. Authenticity of yeast strains was confirmed by polymerase chain reaction (PCR)-based genotyping. Yeast cells were cultured in either YPD (1% yeast extract, 2% peptone, and 2% dextrose [w/v]) or YPGE (3% glycerol + 1% ethanol [w/v]) medium. Solid YPD and YPGE media were prepared by addition of 2% (w/v) agar. For metal supplementation experiments, growth medium was supplemented with divalent chloride salts of Cu, Mn, Mg, Zn or FeSO_4_. For growth on solid media, 3 μl of 10-fold serial dilutions of pre-cultures were seeded onto YPD or YPGE plates and incubated at 37°C for the indicated period. For growth in the liquid medium, yeast cells were pre-cultured in YPD and inoculated into YPGE and grown to mid-log phase. To acidify or alkalinize liquid YPGE, equivalents of HCl or NaOH were added, respectively. Liquid growth assays in acidified or alkalinized YPGE, cultures were grown for 42 h before comparing growth. For growth in the presence of concanamycin A (ConcA), cells were first cultured in YPD, transferred to YPGE allowed to grow for 24 h, then ConcA was added and allowed to grow further for 20 h. Growth in liquid media was monitored spectrophotometrically at 600 nm.

**Table 1:** *Saccharomyces cerevisiae* strains used in this study.

| Yeast Strains | Genotype | Source |
| --- | --- | --- |
| BY4741 WT | *MATa, his301, leu200, met1500, ura300* | Greenberg, M.L. |
| BY4741 *coa6Δ* | *MATa, his301, leu200, met1500, ura300, coa6Δ::kanMX4* | Open Biosystems |
| BY4741 *gef1Δ* | *MATa, his301, leu200, met1500, ura300, gef1Δ::kanMX4* | Open Biosystems |
| BY4741 *aps3Δ* | *MATa, his301, leu200, met1500, ura300, aps3Δ::kanMX4* | Open Biosystems |
| BY4741 *aps3Δ* - NMG | *MATa, his301, leu200, met1500, ura300, aps3Δ::kanMX4* | This study |
| BY4741 *apm3Δ* | *MATa, his301, leu200, met1500, ura300, apm3Δ::kanMX4* | Open Biosystems |
| BY4741 *apl5Δ* | *MATa, his301, leu200, met1500, ura300, apl5Δ::kanMX4* | Open Biosystems |
| BY4741 *apl6Δ* | *MATa, his301, leu200, met1500, ura300, apl6Δ::kanMX4* | Open Biosystems |
| BY4741 *rim20Δ* | *MATa, his301, leu200, met1500, ura300, rim20Δ::kanMX4* | Open Biosystems |
| BY4741 *rim20Δ* - NMG | *MATa, his301, leu200, met1500, ura300, rim20Δ::kanMX4* | This study |
| BY4741 *rim21Δ* | *MATa, his301, leu200, met1500, ura300, rim21Δ::kanMX4* | Open Biosystems |

### Construction of yeast deletion pool

The yeast deletion collection for Bar-Seq analysis was derived from the Variomics library constructed previously (26) and was a kind gift of Xuewen Pan. The heterozygous diploid deletion library was sporulated and selected in liquid haploid selection medium (SC-Arg-His-Leu+G418+Canavanine) to obtain haploid cells containing gene deletions. To do this, we followed previously described protocol (26) with the following modification of adding uracil to allow the growth of deletion library lacking *URA3*. Prior to sporulation, the library pool was grown under conditions to first allow loss of *URA3* plasmids and then subsequent selection for cells lacking *URA3* plasmids. Original deletion libraries were initially constructed where each yeast open reading frame was replaced with *kanMX4* cassette containing two gene specific barcode sequence referred to as the UP tag and the DN tag since they are located upstream and downstream of the cassette (75), respectively.

### Pooled growth assays

A stored glycerol stock of the haploid deletion pool containing 1.5 × 10^8^ cells/ml (equivalent of 3.94 optical density/ml) was thawed and approximately 60 μl was used to inoculate 6 ml of YPD, YPGE or YPGE + 5 μM CuCl_2_ media in quadruplicates in 50 ml falcon tube at a starting optical density of 0.04, which corresponded to ∼1.5 × 10^6^ cells/ml. The cells were grown at 30°C in an incubator shaker at 225 rpm till they reached an optical density of ∼5.0 before harvesting. Cells were pelleted by centrifugation at 3000×*g* for 5 min and washed once with sterile water and stored at −80°C. Frozen cell pellets were thawed and resuspended in sterile nanopure water and counted. Genomic DNA was extracted from 5 × 10^7^ cells using YeaStar Genomic DNA kit (Catalog No.D2002) from Zymo Research. The extracted DNA was used as a template to amplify barcode sequence by PCR, followed by purification of amplified DNA by QIAquick PCR purification kit from Qiagen. The number of PCR cycles used for amplification was determined by Quantitative real time PCR such that barcode sequences were not amplified in a nonlinear way. The amplified UP and DN barcode DNA were purified by gel electrophoresis and sequenced on Illumina HiSeq 2500 with 50 base pair, paired-end sequencing at Genomics and Bioinformatics Service of Texas A&M AgriLife Research.

### Assessing fitness of barcoded yeast strains by DNA sequencing

The sequencing reads were aligned to the barcode sequences using Bowtie2 (version 2.2.4) with the -N flag set to 0. Bowtie2 outputs were processed and counted using Samtools (version 1.3.1). Barcode sequences shorter than 15nts or were mapped to multiple reference barcodes were discarded. We noted that the DN tag sequences were missing for many genes and therefore we only used UP tag sequences to calculate the fitness score using T statistics.

### Gene Ontology analysis

To identify enriched gene ontology terms, we generated a rank ordered list based on T-Scores (Supplementary Table 1 and 3) and used the reference genome for *Saccharomyces cerevisiae* in GOrilla (http://cbl-gorilla.cs.technion.ac.il/).

### Cellular and mitochondrial copper measurements

Cellular and mitochondrial copper levels were measured by inductively coupled plasma (ICP) mass spectrometry using NexION 300D instrument from PerkinElmer Inc. Briefly, intact yeast cells were washed twice with ultrapure metal-free water containing 100 μM EDTA (TraceSELECT; Sigma) followed by two more washes with ultrapure water to eliminate EDTA. For mitochondrial samples, the same procedure was performed using 300 mM mannitol (TraceSELECT; Sigma) to maintain mitochondrial integrity. After washing, samples were weighed, digested with 40% (w/v) nitric acid (TraceSELECT; Sigma) at 90°C for 18 h, followed by 6 h digestion with 0.75% H_2_O_2_ (Sigma-Supelco), then diluted in ultrapure water and analyzed. Copper standard solutions were prepared by diluting commercially available mixed metal standards (BDH Aristar Plus).

### Subcellular fractionation

Whole-cell lysates were prepared by resuspending ∼100 mg of yeast cells in 350 μl SUMEB buffer (1.0% [w/v] sodium dodecyl sulfate, 8 M urea, 10 mM MOPS, pH 6.8, 10 mM EDTA, 1 mM Phenylmethanesulfonyl fluoride [PMSF] and 1X EDTA-free protease inhibitor cocktail from Roche) containing 350 mg of acid-washed glass beads (Sigma-Aldrich). Samples were then placed in a bead beater (mini bead beater from Biospec products), which was set at maximum speed. The bead beating protocol involved five rounds, where each round lasted for 50 s followed by 50 s incubation on ice. Lysed cells were kept on ice for 10 min, then heated at 70°C for 10 min. Cell debris and glass beads were spun down at 14,000×*g* for 10 min at 4°C. The supernatant was transferred to a separate tube and was used to perform SDS-PAGE/Western blotting.

Mitochondria were isolated as described previously (76). Briefly, 0.5-2.5 g of cell pellet was incubated in DTT buffer (0.1 M Tris-HCl, pH 9.4, 10 mM DTT) at 30°C for 20 min. The cells were then pelleted by centrifugation at 3,000×*g* for 5 min, resuspended in spheroplasting buffer (1.2 M sorbitol, 20 mM potassium phosphate, pH 7.4) at 7 ml/g and treated with 3 mg zymolyase (US Biological Life Sciences) per gram of cell pellet for 45 min at 30°C. Spheroplasts were pelleted by centrifugation at 3,000×*g* for 5 min then homogenized in homogenization buffer (0.6 M sorbitol, 10 mM Tris-HCl, pH 7.4, 1 mM EDTA, 1 mM PMSF, 0.2% [w/v] bovine serum albumin (BSA) [essentially fatty acid-free, Sigma-Aldrich]) with 15 strokes using a glass teflon homogenizer with pestle B. After two centrifugation steps for 5 min at 1,500×*g* and 4,000×*g*, the final supernatant was centrifuged at 12,000×*g* for 15 min to pellet mitochondria. Mitochondria were resuspended in SEM buffer (250 mM sucrose, 1 mM EDTA, 10 mM MOPS-KOH, pH 7.2, containing 1X protease inhibitor cocktail from Roche).

Isolation of pure vacuoles was performed as previously described (77). Yeast spheroplasts were pelleted at 3,000×*g* at 4°C for 5 min. Dextran-mediated spheroplast lysis of 1 g of yeast cells was performed by gently resuspending the pellet in 2.5 ml of 15% (w/v) Ficoll400 in Ficoll Buffer (10 mM PIPES/KOH, 200 mM sorbitol, pH 6.8, 1 mM PMSF, 1X protease inhibitor cocktail) followed by addition of 200 μl of 0.4 mg/ml dextran in Ficoll buffer. The mixture was incubated on ice for 2 min followed by heating at 30°C for 75 s and returning the samples to ice. A step-Ficoll gradient was constructed on top of the lysate with 3 ml each of 8%, 4%, and 0% (w/v) Ficoll400 in Ficoll Buffer. The step-gradient was centrifuged at 110,000×*g* for 90 min at 4°C. Vacuoles were removed from the 0%/4% Ficoll interface.

Crude cytosolic fractions used to quantify Sod1 activity and abundance were isolated as described previously (78). Briefly, ∼70 mg of yeast cells were resuspended in 100 μl of solubilization buffer (20 mM potassium phosphate, pH 7.4, 4 mM PMSF, 1 mM EDTA, 1X protease inhibitor cocktail, 1% [w/v] Triton X-100) for 10 min on ice. The lysate was extracted by centrifugation at 21,000×*g* for 15 min at 4°C, to remove the insoluble fraction. Protein concentrations for all cellular fractions were determined by the BCA assay (Thermo Scientific).

### SDS-PAGE and Western blotting

For SDS-polyacrylamide gel electrophoresis (SDS-PAGE)/Western blotting experiments, 20 μg of protein was loaded for either whole cell lysate or mitochondrial samples, while 30 μg of protein was used for cytosolic and vacuolar fractions. Proteins were separated on either 4-20% stain-free gels (Bio-Rad) or 12% NuPAGE Bis-Tris mini protein gels (ThermoFisher Scientific) and blotted onto a polyvinylidene difluoride membranes. Membranes were blocked for 1 h in 5% (w/v) nonfat milk dissolved in Tris-buffered saline with 0.1% (w/v) Tween 20 (TBST-milk), followed by overnight incubation with a primary antibody in TBST-milk or TBST-5% (w/v) BSA at 4°C. Primary antibodies were used at the following dilutions: Cox2, 1:50,000 (Abcam 110271); Por1, 1:100,000 (Abcam 110326); Pgk1, 1:50,000 (Life Technologies 459250), Sod1, 1:5,000, and Vma2, 1:10,000 (Sigma H9658). Secondary antibodies (GE Healthcare) were used at 1:5,000 for 1 h at room temperature. Membranes were developed using Western Lightning Plus-ECL (PerkinElmer), or SuperSignal West Femto (ThermoFisher Scientific).

### Enzymatic activities

To measure Sod1 activity, we used an in-gel assay as described previously, (79). Twenty-five μg of cytosolic protein was diluted in NativePAGE sample buffer (ThermoFisher Scientific) and separated onto a 4-16% NativePAGE gel (ThermoFisher Scientific) at 4°C. The gel was then stained with 0.025% (w/v) nitroblue tetrazolium, 0.010% (w/v) riboflavin for 20 min in the dark. This solution was then replaced by 1% (w/v) tetramethylethylenediamine for 20 min and developed under a bright light. The gel was imaged by Bio-Rad ChemiDoc™ MP Imaging System and densitometric analysis was performed using Image Lab software.

CcO and citrate synthase enzymatic activities were measured as described previously (80) using a BioTek’s Synergy™ Mx Microplate Reader in a clear 96 well plate (Falcon). To measure CcO activity, 15 *µ*g of mitochondria were resuspended in 115 *µ*l of CcO buffer (250 mM sucrose, 10 mM potassium phosphate, pH 6.5, 1 mg/ml BSA) and allowed to incubate for 5 min. The reaction was started by the addition of 60 *µ*l of 200 μM oxidized cytochrome *c* (equine heart, Sigma) and 25.5 *µ*l of 1% (w/v) N-Dodecyl-Beta-D-Maltoside. Oxidation of cytochrome *c* was monitored at 550 nm for 3 min, then the reaction was inhibited by the addition of 7 *µ*l of 7 mM KCN. To measure citrate synthase activity, 10 µg of mitochondria were resuspended in 100 *µ*l of citrate synthase buffer (10 mM Tris-HCl pH 7.5, 0.2% [w/v] Triton X-100, 200 *µ*M 5,5’-dithio-bis-[2-nitrobenzoic acid]) and 50 *µ*l of 2 mM acetyl-CoA and incubated for 5 min. To start the reaction, 50 *µ*l of 2 mM oxaloacetate was added and turn-over of acetyl-CoA was monitored at 412 nm for 10 min. Enzyme activity was normalized to that of WT for each replicate.

### Measuring vacuolar pH

Vacuolar pH was measured using a ratiometric pH indicator dye, BCECF-AM (2′,7′-bis-(2-carboxyethyl)-5-(and-6)-carboxyfluorescein [Life Technologies]) as described by (81) using a BioTek’s Synergy™ Mx Microplate Reader. Briefly, 100 mg of cells were resuspended in 100 *µ*l of YPGE containing 50 *µ*M BCECF-AM for 30 min shaking at 30°C. To remove extracellular BCECF-AM, cells were washed twice and resuspended in 100 *µ*l of fresh YPGE. 25 *µ*l of this cell culture was added to 2 ml of 1 mM MES buffer, pH 6.7 or 5.0. The fluorescence emission intensity at 535 nm was monitored by using the excitation wavelengths 450 and 490 nm in a clear bottom black 96 well plate, (Falcon). A calibration curve of the fluorescence intensity in response to pH was carried out as described (81).

### Statistics

T-scores for each pairwise media comparison (e.g. YPD vs. YPGE) were calculated using Welch’s two-sample t-test for yeast knockout barcode abundance values normalized for sample sequencing depth (i.e. counts per million). Statistical analysis on bar charts was conducted using two-sided students *t*-Test. Experiments were performed in three biological replicates, where biological replicates are defined as experiments performed on different days and different starting pre-culture. Error bars represent the standard deviation, *(*P*<0.05), **(*P<*0.01), ***(*P<*0.001).

### Data availability

All data are available in the main text or the supporting supplementary information. Raw sequencing data are available upon request to

## FUNDING AND ADDITIONAL INFORMATION

This work was supported by the National Institutes of Health awards R01GM111672 to VMG, R0GM1097260 to CDK, and 5F31GM128339 to NMG and the Welch Foundation Grant (A-1810) to VMG. NMG was also supported by National Science Foundation award HRD-1502335 in the first year of this work. The content is solely the responsibility of the authors and does not necessarily represent the official views of the National Institutes of Health or the National Science Foundation.

## ACKNOWLEDGEMENTS

We thank Valentina Canedo Pelaez for assistance with growth measurements and Dr. Thomas Meek for allowing us to use the BioTek’s Synergy™ Mx Microplate Reader. We gratefully acknowledge Xuewen Pan for kind gift of the Variomics library from which the barcoded deletion pool used here was derived.

## AUTHOR CONTRIBUTIONS

VMG conceptualized the project. VMG, NMG, and ATG designed the experiments. NMG, ATG, MZ, performed the experiments. CDK and CQ designed the Bar-Seq protocols and generated the yeast deletion collection. NMG, ATG, and VMG analyzed the data and wrote the manuscript. VMG supervised the whole project and was responsible for the resources and primary funding acquisition. All authors commented on the manuscript.

