## Supplementary Figures for "A genome wide copper-sensitized screen identifies novel regulators of mitochondrial cytochrome c oxidase activity"

Table of Contents

Supplementary Figure 1..........S-2
Supplementary Figure 2..........S-3
Supplementary Figure 3..........S-4
Supplementary Figure 4..........S-5
Description of Supporting Data Not Provided in PDF Format..........S-6

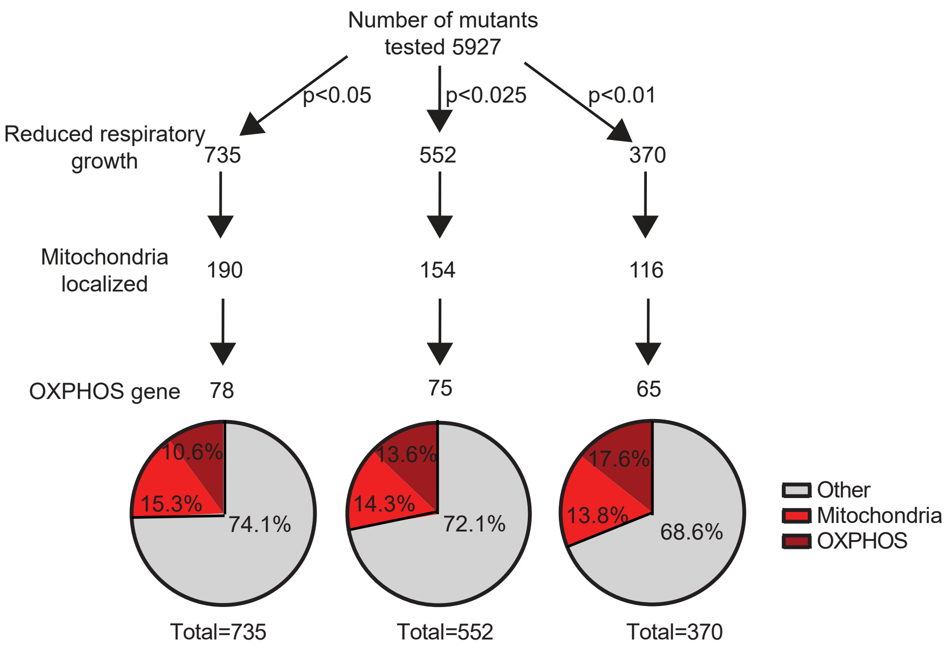


**Supplementary Figure S1.** Enrichment of OXPHOS and other mitochondrial genes increases with increased stringency of p-value threshold.
Illustration of the number of respiratory deficient strains identified at each p-value threshold. The distribution of hits between known OXPHOS protein encoding genes (maroon), other mitochondrial protein encoding genes (red), and non-mitochondrial protein encoding genes (grey) is shown.

### Other genes encoding mitochondrial proteins

#### TCA Cycle
Fum1
Idh1
Kgd1
Kgd2
Mdh1

#### Additional mitochondrial hits
Aat1
Aim17
Aim24
Aim26
Ald4
Apj1
Cym1
Ddl1
Gep7
Gut2
Iai11
Mcp2
Mfm1
Mgr1
Mic60
Mrs2
Nfu1
Num1
Pis1
Puf3
Put1
Rrg7
Sap185
Sod2
Tim21
Uth1
Yhm2
Ysc83

#### Mitochondrial DNA expression
Aim23
Mba1
Mod5
Mrp49
Mrpl1
Mrpl39
Mrx6
Msc6
Oms1
Pet20
Ppe1
Pth2
Pth4
Qri7

### Glutathione biosynthesis
Gsh1
Gsh2

### Rim101 pathway
Dfg16
Rim8
Rim9
Rim13
Rim20
Rim21
Rim101

### AP-3 Complex
Apl5
Apl6
Apm3
Aps3

### Other vesicle mediated transport
Age2
Apl2
Apl4
Apm1
Aps1
Arf2
Arl3
Avl9
Bmh2
Ccz1
Did4
Gcs1
Get1
Get2
Gyp1
Inp53
Irc6
Mon1
Pep8
Ras2
Ric1
Rtt10
Rvs167
Shr3
Snf8
Svp26
Swa2
Sys1
Tda3
Vab2
Vam7
Vam10
Vps5
Vps17
Vps29

### Other vesicle mediated transport continued
Vps35
Vps41
Vps51
Vps63
Yck3
Yip4
Ypt7

### Ion and other transport
Ant1
Atx1
Avt5
Ccc1
Erc1
Flc1
Flc2
Flc3
Fre7
Ftr1
Gdt1
Gef1
Hxt8
Kha1
Mch5
Nhx1
Osh3
Rav1
Rav2
Sam3
Stl1
Stv1
Vtc5
YDL199C
YGR125W
YOL162W

### Other hits

| | | | |
|---|---|---|---|
| Aht1 | Jnm1 | Rim11 | YDR048C |
| Adh6 | Kns1 | Rmd1 | YDR269C |
| Apt2 | Ktr2 | Rmd8 | YDR271C |
| Arg80 | Ldb18 | Rot2 | YDR391C |
| Asi3 | Lug1 | Rpl16B | YDR455C |
| Azf1 | Mak10 | Rrd2 | YER097W |
| Ber1 | Mbr1 | Rtc1 | YGL199C |
| Bmh1 | Mdh2 | Rvs167 | YGL235W |
| Bye1 | Mds3 | Sas3 | Ygp1 |
| Car1 | Met16 | Scs22 | YHL005C |
| Car2 | Mft1 | Sds3 | YJR087W |
| Cat8 | Mnl2 | Ser3 | YJR120W |
| Cnn1 | Mrk1 | Sgn1 | YKL136W |
| Cti6 | Mth1 | Ski7 | YLL020C |
| Ctt1 | Msc6 | Sko1 | YLR169W |
| Cub1 | Msn1 | Smi1 | YLR202C |
| Cys4 | Nab6 | Snf3 | YLR269C |
| Dat1 | Nbp2 | Snf4 | YLR345W |
| Dbr1 | Ndl1 | Sok1 | YML082W |
| Dcs1 | Ngl2 | Sok2 | YML089C |
| Ddr48 | Nis1 | Spg3 | YML090W |
| Eaf7 | Npr1 | Spo7 | YMR057C |
| Edc3 | Npr2 | Ssa1 | YMR315W |
| Erv2 | Nsi1 | Ste20 | Yng1 |
| Far10 | Ntg2 | Syh1 | Ynk1 |
| Fbp1 | Nto1 | Toh1 | YNL140C |
| Fbp26 | Nut1 | Tor1 | YNL179C |
| Fsh1 | Opi1 | Trm10 | YNR042W |
| Fyv8 | Orm2 | Ubr1 | YOL036W |
| Gal11 | Par32 | Ume1 | YOR139C |
| Gal7 | Pcl6 | Ura4 | YOR333C |
| Gal80 | Pcl8 | Vba4 | YPL150W |
| Gal83 | Pho2 | Vhs1 | YPL182C |
| Gdh1 | Pho4 | Vms1 | YPR012W |
| Glc8 | Pph21 | Vps61 | YPR039W |
| Gzf3 | Ppm1 | YAL004W | YPR050C |
| Hap4 | Ppq1 | Yap5 | YPR114W |
| Hap5 | Psp2 | YBL044W | YPR123C |
| Hda2 | Psy2 | YBL055C | Yps7 |
| Hda3 | Ptk1 | YBL096C | Yta7 |
| Hel2 | Ptp3 | YBR224W | |
| Hir1 | Pub1 | YBR225W | |
| Hir2 | Pus1 | YBR226C | |
| Irc14 | Pyc1 | YDL068W | |
| Isw2 | Rck2 | YDL172C | |

**Supplementary Figure S2.** Respiratory deficient mutants of non-OXPHOS related proteins. Non-OXPHOS associated proteins with a p-value ≤0.01 for T-score (YPGE-YPD) were grouped into different functional categories.

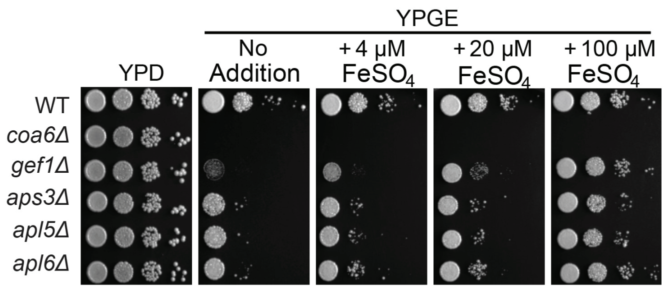


**Supplemental Figure S3.** Respiratory growth of AP-3 mutants is restored by high iron supplementation. Serial dilutions of WT and the indicated mutants were seeded onto YPD and YPGE plates with and without 4, 20 or 100 μM $FeSO_4$ at 37°C and grown for two (YPD) or four days (YPGE). Here *gef1Δ* cells were used as a positive control for iron-based rescue.

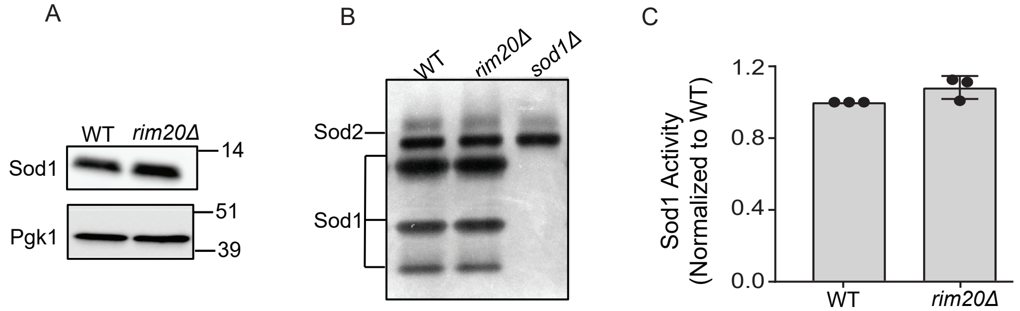


**Supplementary Figure S4.** Activity of cytosolic cuproenzyme Sod1 is not altered in *rim20Δ* cells. (A) Crude cytosolic fraction was analyzed by SDSPAGE/western blot to detect Sod1 abundance in the indicated strains. Pgk1 was used as a loading control. (B) Sod activity in isolated crude cytosolic fraction from the indicated strains was measured by in-gel assay as described in Methods. (C) For quantification of the Sod1 activity in (B), the images were digitalized and densitometry analysis was performed using the Image Lab software.

**Description of Supporting Data Not Provided in PDF Format**

Table S1: Excel file containing a rank ordered list of mutants' growth in YPGE compared to YPD.
Table S2: Excel file containing a list of yeast genes encoding OXPHOS proteins and assembly factors.
Table S3: Excel file containing a rank ordered list of mutants' growth in YPGE+Cu compared to YPGE.
